# MCA1 mechanosensitive channels enable fast communication of wound signals between lateral roots

**DOI:** 10.64898/2025.12.08.692898

**Authors:** Angel Baudon, Shouguang Huang, Dirk Becker, Dietmar Geiger, M. Rob G. Roelfsema, Rainer Hedrich

**Affiliations:** Molecular Plant Physiology and Biophysics, Julius-von-Sachs Institute for Biosciences, Biocenter, University of Würzburg, Julius-von-Sachs-Platz 2, 97082 Würzburg, Germany; Faculty of Synthetic Biology, Shenzhen University of Advanced Technology, Shenzhen, China; Institute of Emerging Agricultural Technology, Shenzhen University of Advanced Technology, Shenzhen, China; State Key Laboratory of Quantitative Synthetic Biology, Shenzhen Institute of Synthetic Biology, Shenzhen Institutes of Advanced Technology, Chinese Academy of Sciences, Shenzhen, China

## Abstract

Although plant roots are hidden in soil, they are vulnerable to damage by insect herbivory, just as like aerial tissues. Yet wound signalling in roots is poorly understood. Here, we examined how damage signals spread locally and over long distances between *Arabidopsis* lateral roots. Using intracellular membrane potential recordings, calcium imaging, and optogenetics, we show that mechanical injury triggers immediate local membrane depolarization and cytosolic Ca^2+^ elevations whose magnitude and duration scale with wound severity. Depolarizations were also detected in neighbouring lateral roots within milliseconds, demonstrating the presence of a rapid inter-root signalling pathway. Through mutant analyses, we highlight the roles of glutamate-like receptors and MCA1 mechanosensitive channels in mediating this long-distance communication. Our results demonstrate that a wound-induced decrease in root cell turgor pressure rapidly spreads across the root network, where neighboring roots decode this signal via MCA1. This work underscores fundamental differences between root and shoot wound responses and uncovers a mechanosensory basis for fast communication between lateral roots.

## Introduction

Throughout evolution, plants have developed defence mechanisms to resist attacks by phytophagous animals, both above and below ground **(Blossey and Hunt-Joshi, 2003)**. Although the root surface represents a crucial plant-to-environment interface and root damage impairs crop production, only a few studies have investigated root defence mechanisms. As a result, our knowledge of plant root defence systems is sparse, and general concepts found in the shoot are often not transposable to roots **(Chuberre et al., 2018)**. Therefore filling this gap, and understanding how plant roots detect and respond to damage caused by pathogens, such as nematodes and insects, is crucial for optimizing crop production.

Apart from bacteria, fungi, and nematodes that can cause single-cell root wounding, insects and their larvae can mechanically damage large areas of roots. Neighbouring root cells can detect the resulting cell debris as Damage-Associated Molecular Patterns (DAMPs), and activate defence mechanisms to protect themselves, such as the production of inducible physical and chemical barriers, and toxic metabolites **(Üstüner et al., 2022)**.

Besides the immediate local defence response at the wounded site, plants also show systemic responses that require communication within or between organs. Chemical or physical messengers can mediate such long-distance signalling. Many studies investigating plant stress signals in the shoot highlighted the importance of fast-propagating electrical signals and intercellular Ca^2+^ waves. For instance, caterpillar feeding on Arabidopsis leaves provokes long-distance leaf-to-leaf Ca^2+^ and electrical waves **(Nguyen et al., 2018; Salvador-Recatalà et al., 2014; Toyota et al., 2018)**. In shoot tissues, several molecules have been identified as contributors to long-distance signalling, including glutamate **(Toyota et al., 2018)** and the β-thioglucoside glucohydrolase **(Gao et al., 2023)**. In addition, physical signals, like variations in turgor pressure, can act as a physical signal transmitted between leaves, with mechanosensors functioning as its decoders **(Moe-Lange et al., 2021)**.

In contrast to systemic signalling in shoots, the mechanisms underlying the initiation and propagation of Ca^2+^ electrical signals within the root system have received little attention so far. In the present study, we addressed this issue and showed that lateral root wounding provokes an immediate depolarization of epidermal cells, followed by a Ca^2+^ signal that spreads locally. At the same time, wounding also triggers depolarization of distant roots that involves glutamate receptor-like cation channels, and mechanosensitive channels. Our results suggest that the drop in pressure due to membrane disruption reaches distant sites within the root and opens mechanosensitive channels to generate a depolarization.

## Results

### Lateral root wounding provokes epidermal cell depolarization and cytosolic Ca^2+^ signals

The elongation zone (EZ) of lateral roots represents a very vulnerable tissue, as cells within this fast-growing part of the root system are not yet protected by a secondary cell wall **(Chuberre et al., 2018)**. The electrical wound responses of these cells were investigated in 10- to 14-day-old seedlings. Single-cell micromanipulator-assisted mechanical wounding of the lateral root tip was performed with a <0.1 µm sharp glass micrcapillary (**Fig.1A**). Wounding a single cell in the root tip triggered an immediate depolarization of 49.4 ± 4.7 mV with a very fast rise (0.78 ± 0.11 s), and a decay of 21.5 ± 6.8 s (n=12, **Fig.1B-C**). In contrast, damaging entire cross-section of the lateral root tip via a broken 10 µm glass capillary resulted in an approximately two times bigger depolarization (91.4 ± 6.3 mV, p<0.0001) with a similar rise time (1.00 ± 0.12 s, p = 0.1921), but a much slower decay time (118.4 ± 21.2 s, p <0.0001, n = 7, **Fig.1B-C**).

**Figure 1.**
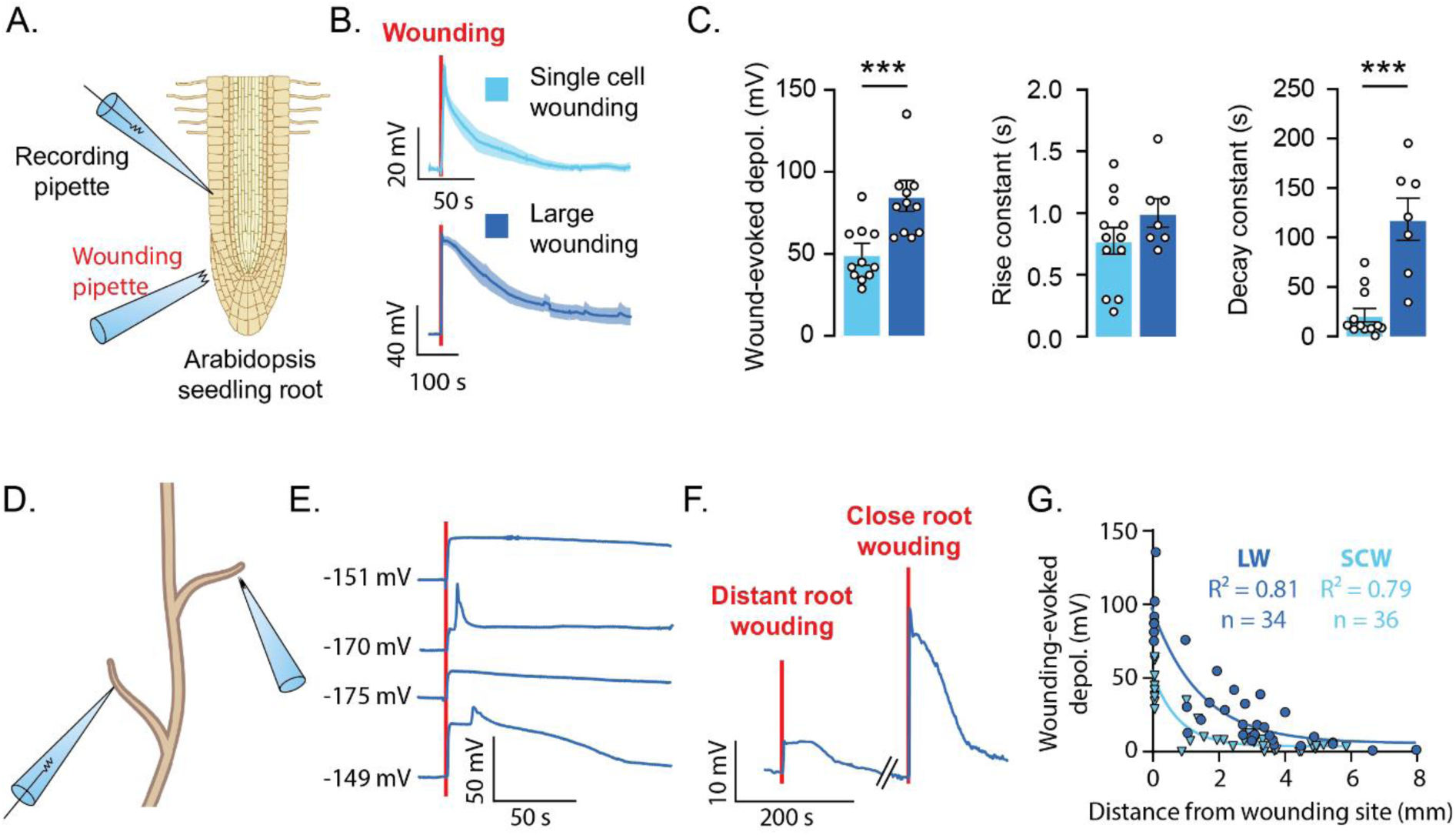
Electrical signals evoked by lateral root wounding. **A.** Lateral root epidermal cells were impaled with a sharp glass pipette to measure the plasma membrane potential, and a single-cell wounding was performed using another sharp glass pipette, or a large wound was made by piercing the entire lateral root with a ∼10µm broken pipette. **B.** Mean depolarization of the membrane potential caused by single-cell wounding (SCW, light blue) or large wounding (LW, dark blue). The vertical red line marks the time point of wounding. **C.** Quantification of the depolarization amplitude, rise time, and decay time caused by single-cell wounding (light blue) or large wounding (dark blue). n_SCW_ = 12, n_LW_ = 7. **D.** An epidermal cell of a lateral root was impaled with a sharp glass pipette, and a large wound was inflicted on another, distant lateral root using a ∼10µm broken pipette. **E.** Example traces showing membrane potential depolarization caused by large wounding. Each trace is from a different plant; the membrane potential at the start of the experiment is given for each trace, and the red line indicates the time point of wounding. **F.** Membrane potential trace showing depolarization triggered by wounding a distant and a nearby lateral root, the trace is interrupted by a period of ∼600s, as indicated by the diagonal lines. **G.** The depolarization amplitude evoked by a single cell wounding (SCW), or a large wounding (LW) plotted against the distance between the wounded root and the recording site. Data are shown as mean across plants ± SEM. Detailed statistics are available in the *Statistics Table*. See also Figure S1. ***p<0.001

To test whether wound-evoked electrical signals spread, we wounded a lateral root while measuring the EZ epidermal cell membrane potential of a neighbouring lateral root (**Fig.1D**). Upon wounding, we detected an almost immediate (millisecond range) depolarization at the distant site in all our recordings (**Fig.1E**). The amplitudes of distant wounding-evoked depolarizations were quite variable and appeared to decrease with the distance between the recording and the wounding site. To resolve the amplitude-distance relation, we impaled a lateral root and performed the consecutive wounding of two different neighbouring lateral roots, one more distant from the recorded root, and another one closer to the recording site. The depolarization observed following a distant root wounding was considerably smaller compared to the one caused by wounding of a closer lateral root (**Fig.1F**), suggesting that the depolarization amplitude depends on the distance.

On average, the wound-evoked depolarization amplitude decayed exponentially with the distance between the wounding and the recording site (**Fig.1G**). The single-cell wounding-evoked depolarization could still be detected up to 2 mm away from the wounding site and had a half-maximum distance of 0.55 mm (**Fig.1G**). When increasing the size of the wound, we could detect the evoked depolarization up to 5mm away from the wounding site (half-maximum distance = 1 mm, **Fig.1G**).

### Wounding evokes Ca^2+^ signalling in roots

Since electrical signals and cytosolic Ca^2+^ elevations often occur together following leaf wounding **(Moe-Lange et al., 2021; Nguyen et al., 2018)**, we investigated whether this is the case in root tissues. Cytosolic Ca^2+^ signals in lateral roots were monitored with the genetically encoded Ca^2+^ indicator GCaMP3 expressed under the 35S promoter. Fluorescence imaging of GCaMP3-expressing seedlings showed that lateral root single cell wounding triggered a 391 ± 27% fluorescence increase that reached a peak within 12 ± 2 s and decayed in 182 ± 50 s (**Fig.2A-C**). Interestingly, increasing the wounding size resulted in a Ca^2+^ transient with a similar amplitude (361 ± 28%, p = 0.4521), and a similar rise time (22 ± 7s, p = 0.2606), but a slower decay (391 ± 27s, p = 0.0023, **Fig.S2A-B**). This response suggests that the duration, rather than the amplitude, of the Ca^2+^ transient reports the dimension of the injury.

**Figure 2.**
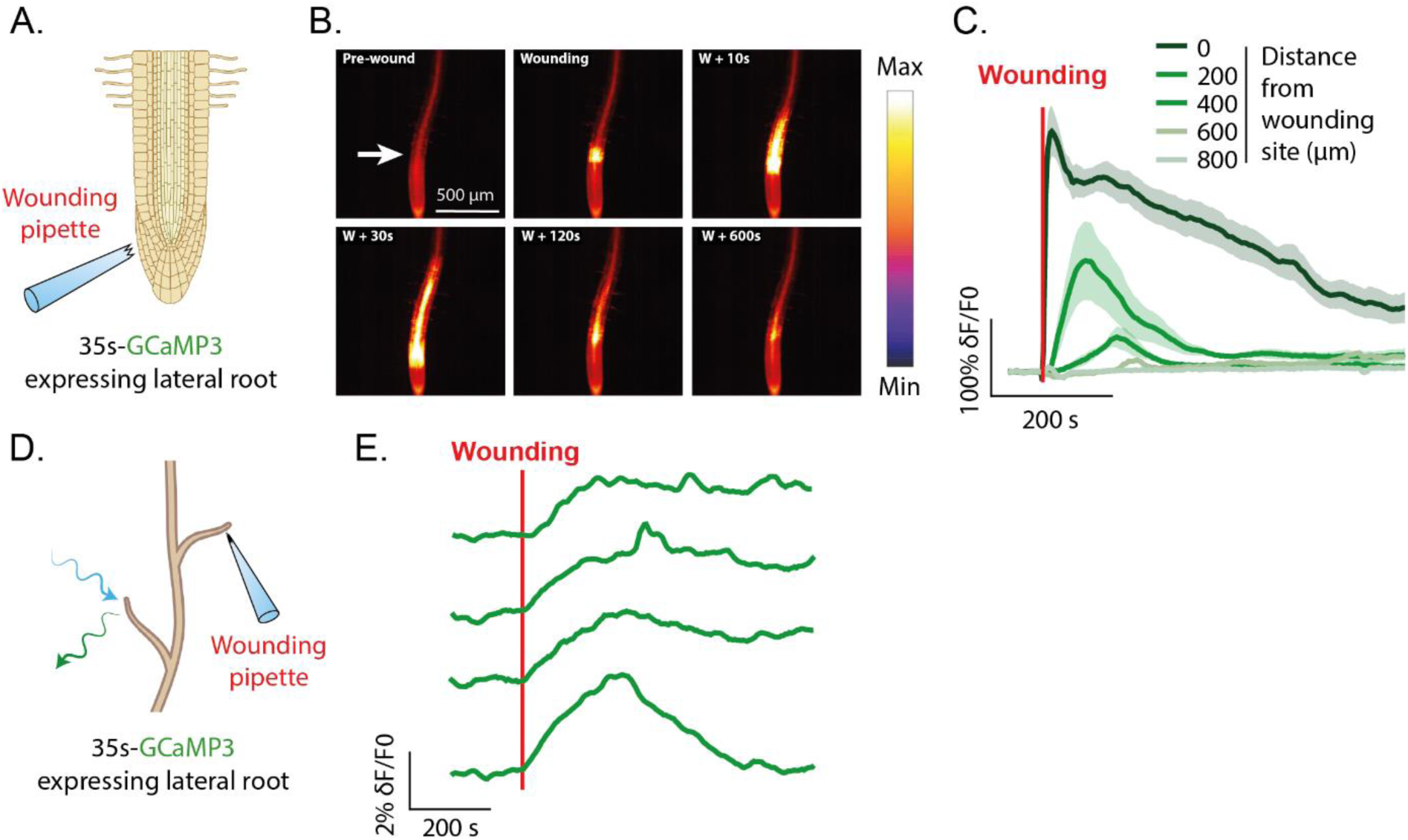
Lateral root wounding-evoked Ca^2+^ signals. **A.** GCaMP3-expressing seedlings were stimulated through large wounding with a ∼10µm glass pipette. The white arrow indicates the wound site. **B.** Fluorescence imaging of a lateral root from a GCaMP3-expressing seedling at different times before, during, and after causing a large wound. n=8. Scale: 500µm. **C.** Average fluorescence levels of GCaMP3 measured at several distances from the wound site. The colour code of the traces is given above the graph. **D.** Lateral roots of GCaMP3-expressing seedlings were stimulated by large wounding of a neighbouring root with a ∼10µm glass pipette. **E.** Example traces of GCaMP3 fluorescence intensity before and after wounding a neighbouring lateral root. Each trace is from an individual wounding experiments. The red line indicates the time point of wounding. Data are expressed as mean values ± SEM. Detailed statistics can be found in the *Statistics Table*. See also Figure S2.

We then measured the spread of wounding-evoked Ca^2+^ signal in lateral roots, and observed that a single-cell wounding-evoked Ca^2+^ transient did not propagate very far (half-maximum distance = 91 µm, **Fig.2A-C**). Ca^2+^ signals generated by large wounding propagated further, but also remained relatively close to the wounded area (half-maximum distance = 193.6 µm, **Fig.S2A-B**). However, GCaMP3 measurement in distant root with a higher magnification (**Fig.2D**) allowed the detection of small Ca^2+^ increases in the elongation zones of roots neighbouring the wounded one (**Fig.2E**). These observations are in stark contrast to what has been observed in the shoot, where a strong and non-decremental Ca^2+^ wave propagate from the wounded leaf to interconnected leaves **(Toyota et al., 2018)**.

### Priming Ca^2+^ signals represses wounding-evoked electrical responses

How does a wound signal propagate to neighbouring roots? To investigate this, we assessed whether electrical activity or Ca^2+^ signalling is the key mediator of wound communication between lateral roots. To do so, we used the light-activated Ca^2+^-permeable ChR2-XXM2.0 channel to impose Ca^2+^ transients non-invasively, allowing for the study of the involvement of Ca^2+^ in physiological mechanisms **(Ding et al., 2024; Huang et al., 2024, 2021; Song et al., 2025)**. Since wounding evoked robust Ca^2+^ transients locally, we used ChR2 XXM2.0 to optically impose repetitive Ca^2+^ transients to disturb Ca^2+^-dependent mechanisms while measuring the membrane potential of EZ root cells with intracellular voltage-recording microelectrodes (**Fig.3A**).

**Figure 3.**
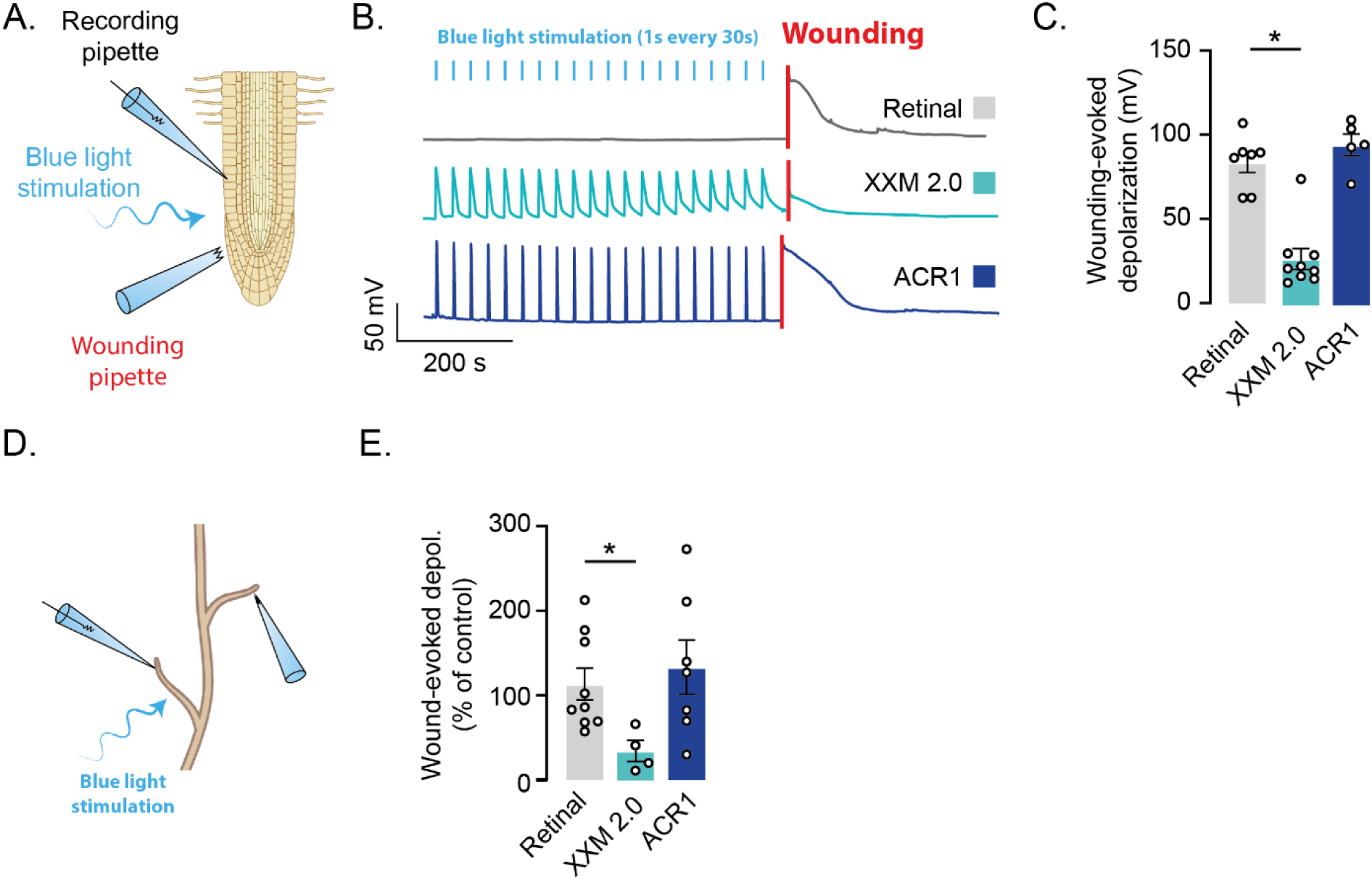
Ca^2+^ involvement in wounding-evoked electrical signalling. **A.** Lateral root epidermal cells were impaled using sharp glass pipettes and stimulated by a light train stimulation consisting of 20 pulses of 1s of blue light (ʎ=470nm, 150µmol.m^-2^.s^-1^) at an interval of 30s. Wounding of the same lateral root was performed ∼30s following the end of the light train stimulations. **B.** Example traces of the membrane potential recorded in seedlings expressing retinal (grey), the anion-permeable channelrhodopsin GtACR1 (blue), or the cation-permeable channelrhodopsin ChR2-XXM (turquoise). The light blue lines indicate the time point of light stimulation, and the red line indicates the time of wounding. **C.** Quantification of the amplitude of the wounding-evoked depolarization following the repetitive stimulation of seedling expressing retinal (grey, n=7) in combination with ChR2-XXM2.0 (turquoise, n=9), or GtACR1 (blue, n=5). **D.** The epidermal cell of a lateral root was impaled and repetitively stimulated with light pulses before wounding another lateral root with a ∼10µm glass pipette. **E.** Amplitude of distant wound-evoked depolarization normalized by its predicted amplitude based on the distance between the recorded site and the wounding point. Retinal (grey, n=9), ChR2-XXM2.0 (turquoise, n=6), or GtACR1 (blue, n=7). Data are expressed as mean across plants ± SEM. Detailed statistics can be found in the *Statistics Table*. See also Figure S3. *p<0.05

Seedlings expressing ChR2 XXM2.0 under the UBQ10 promoter were primed with a train of blue light pulses (1s every 30s, 20 pulses). This light stimulations triggered robust and reproducible depolarizations, which were absent from seedlings expressing only the retinal co-factor without ChR2 XXM2.0 (**Fig.3B**). Following the priming stimulus, wounding the root triggered a smaller membrane potential depolarization in ChR2-XXM2.0-expressing plants compared to the one observed in their retinal-expressing controls (ChR2 XXM2.2: 27.3 ± 6.2, retinal: 83.6 ± 6.0, p = 0.0192, **Fig.3B-C**). This result indicates that Ca^2+^ is involved in local wound-mediated depolarization. However, ChR2 XX2.0 also triggers large depolarization, which can be a confounding factor. To test this possibility, we used the anion channelrhodopsin GtACR1 as a depolarization control **(Zhou et al., 2021)**. In seedlings expressing GtACR1, a train of blue light pulses (1s every 30s, 20 pulses, **Fig.3A**) repeatedly depolarized the membrane potential of epidermal root cells (**Fig.3B-C**). However, this train of depolarisations did not affect the depolarization caused by wounding (ACR1: 94.11 ± 6.46 mV, retinal: 83.6 ± 6.0 mV, p >0.999, **Fig.3B-C**). Given that repetitive Ca^2+^ transients, but not repetitive depolarizations, reduce the amplitude of the local wounding-evoked root depolarization, it indicates that Ca^2+^ signals are important for wound-evoked electrical responses.

To test whether Ca^2+^ also contributes to the depolarization observed in lateral roots adjacent to the wounded one, we recorded the membrane potential of a lateral root and photostimulated it, prior to wounding another lateral root (**Fig.3D**). Wounding a distant lateral root after Ca^2+^ priming via XXM2.0, triggered a smaller depolarization compared to controls (63.4 ± 19.9 % of the control, **Fig.3E**). In contrast, depolarization priming of the local root via light stimulations of ACR1-expressing seedlings did not affect the amplitude of depolarization triggered by distant root wounding (133.3 ± 31.9 % of the control, **Fig.3E**).

This finding indicates that interfering with Ca^2+^-dependent signalling by imposing repetitive Ca^2+^ transients decreases the wounding-associated depolarization, both in the wounded root and in neighbouring lateral roots.

### DAMPs elicit local and long-distance electrical signalling

What is the factor traveling from one lateral root to another? Following wounding, damage-associated molecular patterns (DAMPs) can be released from wounded cells and convey the wounding information **(Vega-Muñoz et al., 2020)**. To test this hypothesis, we locally applied DAMPs such as glutamate, adenosine triphosphate (ATP), or reactive oxygen species (ROS) using glass pipettes connected to a pressure-ejection system and monitored the voltage response of EZ cells from lateral roots (**Fig.4A**). Locally, all three DAMPs elicited robust depolarizations (**Fig.4B-D**) with similar EC_50_ values (Glutamate: 4.7 mM, ATP: 1.6 mM, H_2_O_2_: 3.1 mM). However, the rise kinetics of the depolarization were different, with glutamate triggering a faster membrane potential increase of epidermal root cells, compared to the other DAMPs (Glutamate: 22.1 ± 4.8 s, ATP: 104.9 ± 24.9 s, H_2_O_2_: 99.67 ± 19.74 s, p = 0.0048, **Fig.4B-D**).

**Figure 4.**
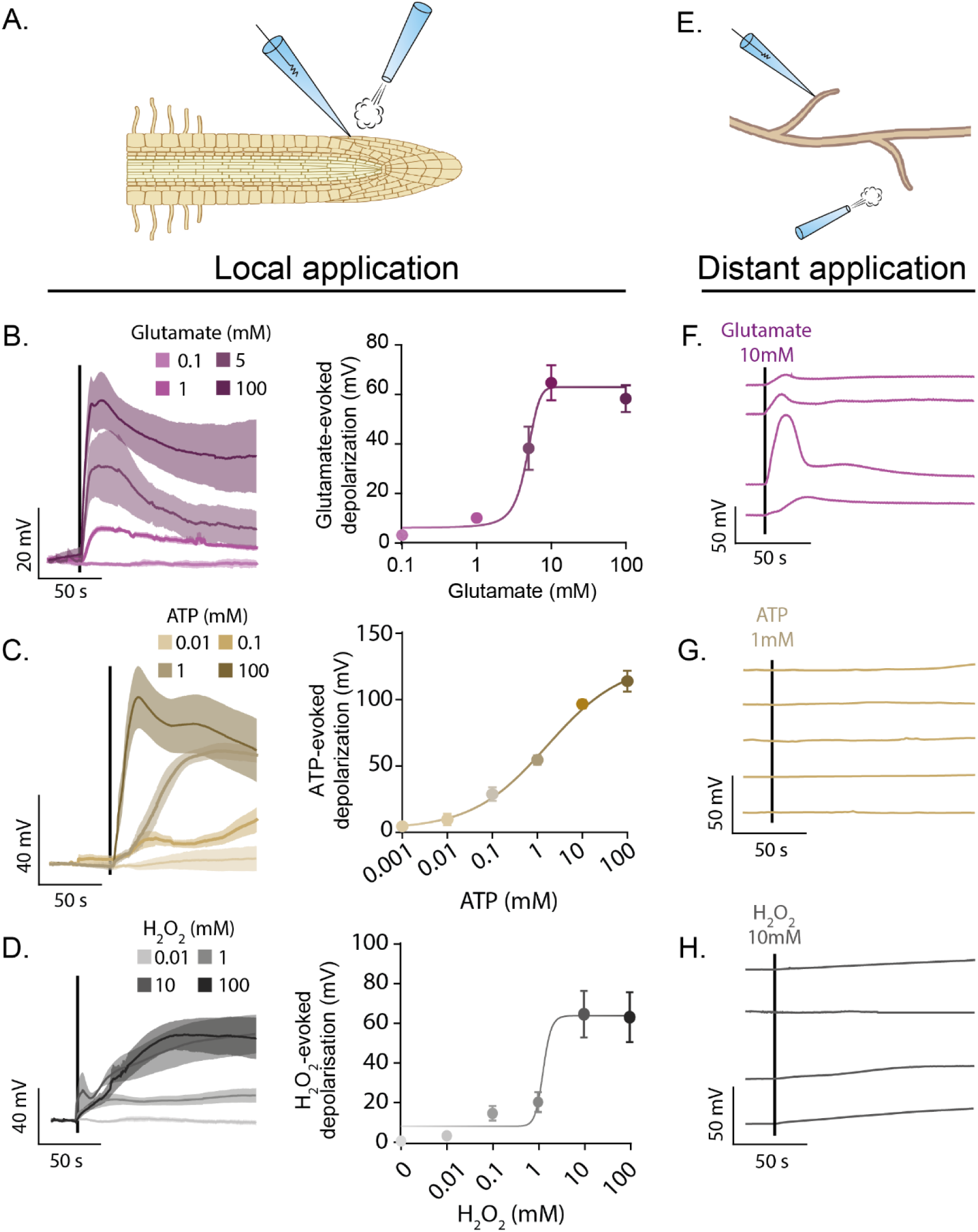
Effect of Damage-Associated Molecular Patterns (DAMPs) on root membrane potential. **A.** Epidermal root cells of lateral roots were impaled, and a solution containing different concentrations of glutamate (**B**), ATP (**C**), or H_2_O_2_ (**D**) was backpressure-applied locally with a second glass pipette for 10s. Mean traces of evoked depolarization at different concentration of DAMPs are shown on the left, and their corresponding quantification on the right. **E.** Epidermal root cells of lateral roots were impaled, and a solution containing glutamate (**F**), ATP (**G**), or H_2_O_2_ (**H**) was backpressure-applied on a neighbouring root with a second glass pipette for 10s. Traces shows membrane potential of epidermal root cells during a distant puff application. Each trace results from an independent recording. Black lines indicates puff applications. Data are expressed as mean across plants ± SEM. Detailed statistics can be found in the *Statistics Table*. See also Figure S4.

To test if the DAMP-mediated depolarizations can travel from a lateral root to another, the membrane potentials of root cells were recorded while DAMPs were locally applied on a neighbouring lateral root (**Fig.4E**). Under these conditions, neither H_2_O_2_ nor ATP were able to generate membrane potential depolarizations in the distant lateral root. Glutamate however, triggered depolarizations in distant lateral roots (**Fig.4F-H**). This observation points toward an important role of glutamate in mediating wound-evoked communication between lateral roots.

### Glutamate-like receptors participate in long-distance wound signalling in roots

Glutamate-like receptors (GLRs), and especially GLR3.3 and GLR3.6, represent potential candidates to detect glutamate, since they are involved in leaf-to-leaf propagation of wounding-evoked Ca^2+^ and electrical signals **(Mousavi et al., 2013; Nguyen et al., 2018)**. To test if these GLRs are involved in wounding-evoked Ca^2+^ and electrical signals in the root, we impaled root epidermal cells of seedlings lacking functional GLR3.3 and GLR3.6 (*glr3.3/3.6*). Local depolarization evoked by glutamate application was reduced in roots of *glr3.3/3.6* mutant plants (**Fig.5A-C**, p > 0.001). To test the involvement of these GLRs in long-distance propagation of glutamate-evoked depolarization, we impaled epidermal cells of a lateral root and applied glutamate to a distant lateral root. In these GLR3.3/3.6-lacking plants, application of glutamate on a distant root only triggered depolarizations of low amplitude (**Fig.S5A**). Altogether, these data make GLR potential candidates for conveying wounding-associated DAMP-mediated depolarization in the root.

**Figure 5.**
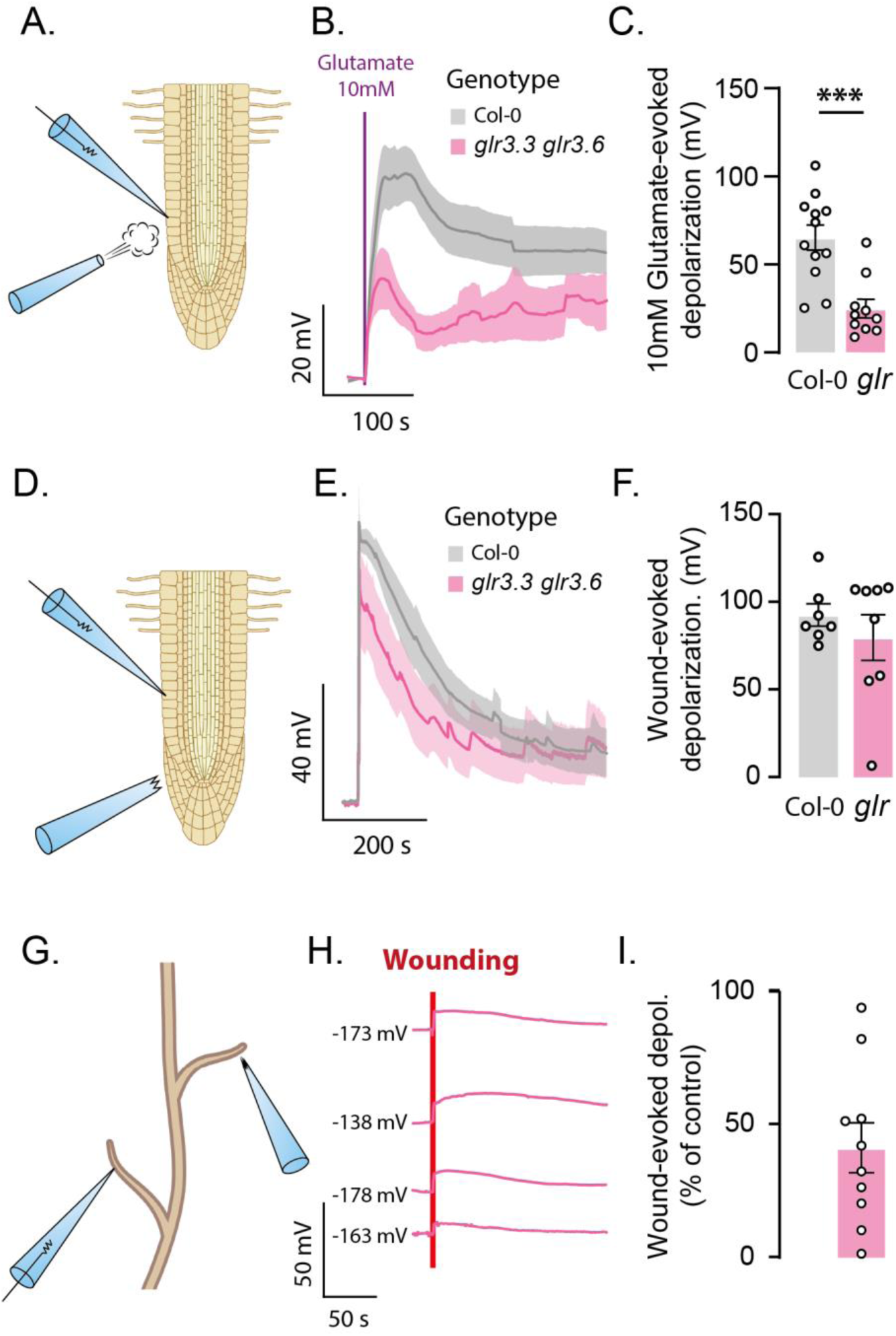
Glutamate-like receptors are important for long-distance wound-evoked depolarizations. **A.** Epidermal root cells were impaled and a solution containing 10mM of glutamate was backpressure-applied locally with a second glass pipette for 10s. **B.** Mean traces of the glutamate-evoked depolarization. The time point of glutamate application is indicated with the purple line, the grey traces come from wild type plants, and the pink ones from *glr3.3/3.6* mutant plants. **C.** Amplitude of glutamate-evoked depolarization. n_Col-0_ = 12, n_glr_ = 10 plants. **D.** Lateral root epidermal cells were impaled, and large wounds were made. **E.** Mean depolarization of the membrane potential evoked by a large wounding in wild type (grey), or *glr3.3/3.6* mutant (pink). The red line indicates the time point of wounding. **F.** Quantification of the wound-evoked depolarization amplitude, rise, and decay constants. n_Col-0_ = 7, n_glr_ = 8 plants. **G.** The epidermal cell of a lateral root was impaled using a sharp electrode, and a large wounding was performed on another, distant lateral root. **H.** Example traces of membrane potential depolarization evoked by a large wounding of *glr3.3/3.6* mutant distant lateral root. Each trace comes from a different plant, the number before the trace indicates the resting membrane potential, and the red line indicates the time point of wounding. **I.** Amplitude of distant wound-evoked depolarization recorded in *glr3.3 glr3.6* mutant plants normalized by their predicted amplitude based on the distance between the recorded site and the wounding point. n = 10. Data are expressed as mean across plants ± SEM. Detailed statistics can be found in the *Statistics Table*. See also Figure S5. ***p<0.001

Given that wounding is a complex event that likely involves different DAMPs, we wondered if impairing the glutamatergic system in the root was enough to prevent wound signalling. In *glr3.3/3.6* mutant plants, the amplitude and kinetics of local wound-evoked depolarization were similar to those observed in Col-0 control plants (p_amplitude_ > 0.999, **Fig.5E-F**, p_rise_ = 0.795, p_decay_ = 0.235, **Fig.S5B**). This behaviour suggests that GLR3.3 and GL3.6 are not key in the generation of the local depolarization evoked by wounding, as shown before **(Marhavý et al., 2019)**. What about the distal sites? To answer this question, we tested if the selected GLRs play a major role in wounding-evoked depolarization at sites distal to the wounding site (**Fig.5G**). Following wounding, we observed small depolarizations of the membrane potential in *glr3.3/3.6* mutant plants (**Fig.5H**). To evaluate whether they were smaller than in wild type plants, we inferred their expected amplitude from their distance using the exponential relationship in Figure 1J. This analysis indicated that wounding-evoked depolarizations observed in *glr3.3/3.6* mutants were significantly smaller (-55.7 ± 8.6 %, **Fig.5I**) compared to those expected in wild type for the same distance (p = 0.002).

Together, these finding indicates that GLRs are involved in the propagation of electric signals from the site of wounding to neighbouring lateral roots.

### Wound-evoked hydrostatic pressure drop activates mechanosensitive channels

Taking a closer look at distant wound-evoked depolarizations, we observed an extremely short delay between the wounding and the depolarization recorded at the distant recording sites (**Fig.1F-G**). To precisely measure the speed of wounding-evoked electrical signal, we simultaneously impaled two lateral roots and wounded one of them (**Fig.6A**). Based on the delay between the onsets of the wounding-provoked electrical signals, we calculated a mean speed of 74.7 ± 18.2 mm/s (**Fig.6B-C**). In comparison, the speed of the wound-evoked Ca^2+^ wave displayed in Fig.1 was 2.59 x 10^-3^ mm/s and thus about four orders of magnitude slower than the depolarization response. This observation indicates that a signal travels several centimetres per second and generates a wounding-associated depolarization in distant lateral roots.

**Figure 6.**
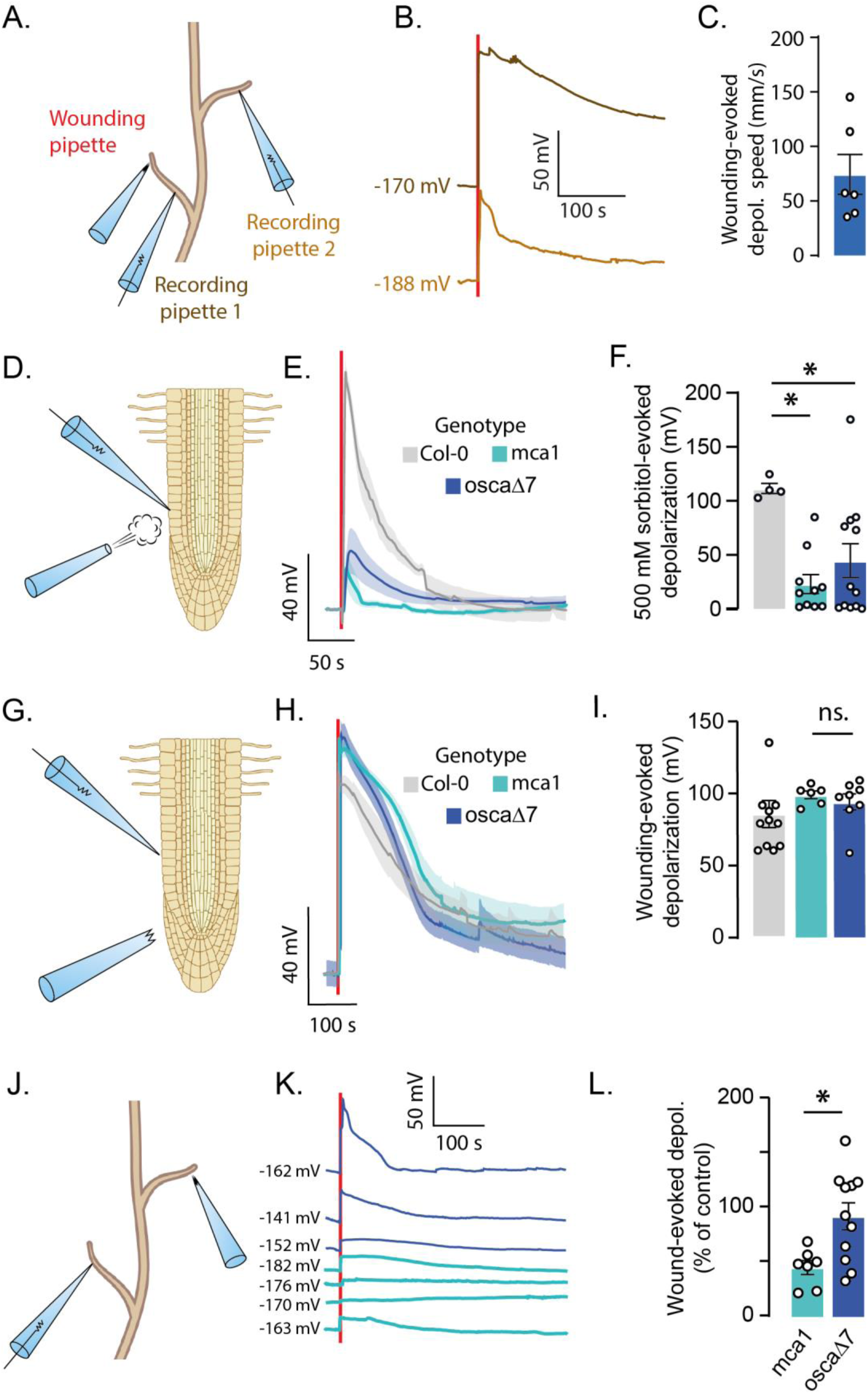
Involvement of mechanosensitive cation channels in wound signalling. **A.** Epidermal cells from two different lateral roots were impaled, and one of them was wounded. **B.** Simultaneously recorded membrane potentials in the wounded root (dark brown, recording pipette 1), and in another lateral root (light brown, recording pipette 2). The red line indicates the time point of wounding. **C.** Propagation speed of the wound-evoked depolarization, calculated based on the time interval between the depolarization onset at the two recording sites, illustrated in A. n=6 plants. **D.** Epidermal root cells were impaled, and a solution containing sorbitol was backpressure-applied locally with a second glass pipette for 1s. **E.** Mean traces of sorbitol-evoked depolarization. Sorbitol application is indicated with the red line. **F.** Amplitude of sorbitol-evoked depolarization. n_Col-0_=4, n_mca1_=10, n_osca1_=12. **G.** Lateral root epidermal cells were impaled, and large wounds were made. **H.** Mean depolarization of the membrane potential evoked by a large wounding in wild type (grey), *mca1* mutant (turquoise), or *osca1* mutant (dark blue). The red line indicates the time point of wounding. **I.** Quantification of the wound-evoked depolarization amplitude. n_Col-0_ = 7, n_mca1_ = 6, n_osca1_ = 8 plants. **J.** The epidermal cell of a lateral root was impaled using a sharp electrode, and a large wound was provoked on another distant lateral root. **K.** Example traces of membrane potential depolarization evoked by a large wounding of *mca1* (turquoise) or *osca1* (dark blue) mutants, distant lateral root. Each trace comes from a different plant, the number before the trace indicates the resting membrane potential, and the red line indicates the wounding. **L.** Amplitude of distant wound-evoked depolarization recorded in *mca1* or *osca1* mutant plants normalized by their predicted amplitude based on the distance between the recorded site and the wounding point. n_mca1_=7, n_osca1_=11. Data are expressed as mean across plants ± SEM. Detailed statistics can be found in the *Statistics Table*. See also Figure S6. *p<0.05.

Which biologically relevant signal can travel that fast? Plant cells represent a hydrodynamic system that is poised to sense changes in hydrostatic pressure. Given that pressure waves travel with the speed of sound, we hypothesize that the sudden loss of turgor pressure due to wounding could propagate to interconnected neighbouring cells and pipes like the xylem/phloem-based vasculature. To investigate whether a local drop of turgor pressure without rupturing the plasma membrane could trigger a depolarization of EZ root cells, we locally applied a hyperosmotic solution of sorbitol for a very short time (1s). When applying 500 mM of sorbitol, the recorded EZ epidermal cells immediately depolarized by 111.5 ± 4.6 mV (**Fig.S6A-C**). Such sorbitol-mediated hyperosmotic shocks were sufficient to trigger local Ca^2+^ signals (**Fig.S6D-I**), which indicates that mechanosensitive elements sense changes in the root cells’ pressure and generate a Ca^2+^ signal.

In plants, several types of mechanosensitive plasma membrane ion channels have been identified, such as MSL-type anion channels, OSCA-type, as well as MCA-type Ca^2+^ conductive cation channels **(Guichard et al., 2022)**. Since Ca^2+^ ions are probably involved in wound-mediated depolarization (**Fig.2-3**), we investigated the role of Ca^2+^-permeable mechanosensitive channels MCA1 and OSCA1 mutants, and recorded membrane potential changes in lateral roots upon local application of a 500 mM sorbitol solution (**Fig.6D**). While local application of sorbitol elicited a strong depolarization in wild type plants (111.5 ± 4.6 mV, n=4), the same challenge provoked a much weaker depolarization in *osca1* and *mca1* mutant plants (*mca1*: 23.1 ± 8.8 mV, n=10, p= 0.011, *osca1*: 44.7 ± 15.7, n=12, p = 0.22, **Fig.6E-F**). Thus, OSCA1 and MCA1 can translate the turgor pressure drop into a membrane depolarization.

During a wounding event, the plasma membrane breaks, and the turgor pressure of the cell is lost. To test if OSCA1 and MCA1 channels are involved in the wounding-induced local depolarization, we monitored the membrane potential of lateral roots. However, no difference in the amplitude and kinetics of the wound-depolarization in *osca1* and *mca1* plants compared to wild type was observed (Col-0: 92 ± 6.0 mV, n=7, mca1: 98.8 ± 2.7 mV, n=6, p = 0.265, osca1: 94.0 ± 15.7, n=8, p = 0.523, **Fig.6G-I**). Thus, although MCA1 and OSCA1 are involved in turgor sensing they do not mediate the depolarization following local wounding.

We hypothesized that a wound-mediated pressure drop sets off a signal that travels from a wounded lateral root to a distal one. This proposed pressure wave could activate mechanosensitive channels that depolarize cells at the distal site. Indeed, after wounding a distant lateral root, we observed a reduction of the depolarization amplitude by approximately 50 % in *mca1*, but not in *osca1* mutant plants compared to the control (**Fig.6J-L**). This indicates that MCA1 channels are involved in decoding the mechanical long-distance signals from the wounding site to neighbouring lateral roots.

## Conclusion & Discussion

In the present study, we studied if and how roots of Arabidopsis seedlings communicate with long-distance electrical and Ca^2+^ signals after wounding. Upon wounding of a lateral root, a Ca^2+^ transient and a depolarization were evoked both locally and in adjacent lateral roots (**Fig.1-2**). Using optogenetic tools, we observed that wounding-evoked depolarization probably relies on Ca^2+^ ions, and modifying its homeostasis impaired wound-mediated depolarization locally (**Fig.3**). By examining the molecular components of the wound-mediated depolarization, we identified glutamate (**Fig.4**), glutamate-like receptors (**Fig.5**), and MCA1 mechanosensitive channels (**Fig.6**) as crucial actors in this process. Together our findings suggest that root wounding provokes a drop in turgor pressure that rapidly propagates to neighbouring lateral roots, activating mechanosensitive MCA1 cation channels and initiating a depolarization, which is amplified by glutamate-like receptors.

This hypothesis implies that the wound-induced depolarization response should scale with the magnitude of the pressure drop. Supporting this, the comparison between single-cell and large-scale wounding reveals that single-cell injury elicits a smaller, shorter-lived, and less far-reaching depolarization (**Fig.1**). These findings are in agreement with Marhavý and colleagues **(Marhavý et al., 2019)**, who used laser ablation to wound a single cortex root cell while measuring extracellular root potentials using surface electrodes. Upon these conditions, they reported that single-cell wounding generates an extracellular potential detectable only within ∼200 µm of the wound site **(Marhavý et al., 2019)**. In our study, intracellular membrane potential recording allowed us to detect cell depolarization up to 2 mm, and large wounding-evoked depolarization, could travel up to 5 mm (**Fig.1**). Although this is sufficient to reach neighbouring lateral roots, this distance is considerably shorter compared to what has been observed in the shoot, where wounding-associated depolarizations can travel several centimetres **(Mousavi et al., 2013; Nguyen et al., 2018)**.

Because roots lie within a substrate that slows insect movement, the local restriction of root wound signalling makes sense; insects that inflict root damage cannot travel through soil as easily as they move above ground. Thus, conveying information about an herbivory-induced attack to distant organs is likely less critical for roots than for shoots. For the same reason, one might also argue that electrical propagation does not need to be as fast as in the shoot. However, we measured a propagation speed that was very similar to leaf-to-leaf signalling speed (tens of mm/s) **(Malone, 1992)**.

Another striking difference of wound-evoked signals between the root and shoot tissue is the ability of seedlings roots to generate a distant wound-evoked depolarization even in the absence of GLR3.3 and GLR3.6. Given that the absence of these channels completely prevents wound-triggered electrical signals from spreading between leaves **(Mousavi et al., 2013; Nguyen et al., 2018)**, and between the root and the shoot **(Shao et al., 2020)**, one might expect a similar lack of long-distance signaling in roots. However, we detected wound-induced depolarizations—albeit with lower amplitude—in lateral roots adjacent to the injured site. This divergence between root and shoot responses indicates a major difference in the channels/mechanisms implicated in long-distance wounding. Recent work has shown that GLRs are inhibited by a Ca²⁺/calmodulin module **(Yan et al., 2024)**. In light of this finding, the desensitization of GLRs induced by optogenetic priming could account for the reduced wound-evoked depolarization observed after repeated Ca²⁺ transients. In line with this hypothesis, and given the residual depolarization observed in *glr3.3/3.6* mutant plants following Ca²⁺ priming, we propose that GLRs amplify the wound signal, rather than initiate it.

The link between Mechanosensitive channels have recently emerged as key components in long-distance wound signalling in leaf-to-leaf communication. While the MSL10 channel appears to play a key role in wound-induced depolarization in the shoot **(Moe-Lange et al., 2021)**, our findings suggest that MCA1 is the primary mechanosensor in the root. The participation of mechanosensitive channels and the observed speed of depolarization suggest that both root and shoot tissues likely rely on a shared wound signaling mechanism triggered by a drop in turgor pressure. However, different molecular players appear to decode this mechanical cue in each tissue.

More broadly, our data aligns with the squeeze cell hypothesis **(Gao and Farmer, 2023)**. According to this hypothesis, wounding induces an axial pressure change that propagates throughout the water column within the xylem. This pressure then spreads radially, to activate mechanosensitive channels, and to generate a GLR-dependent slow wave potential **(Farmer et al., 2014)**. However, it is more probable that wounding the root triggers a drop in turgor pressure, at least when the membrane is disrupted. To reconcile these findings, we could propose that according to the osmoelectric siphon mechanism **(Gao and Farmer, 2023)**, cell depolarization triggers the release of water, thereby increasing the internal pressure of the xylem.

Based on our results, we can further ask what role the long-distance wound signal described above plays in communication between roots. Recent work by Ma and colleagues **(Ma et al., 2025)** showed that root wounding activates the ethylene signalling pathway, which in turn regulates jasmonic acid responses in the wounded root. Notably, this regulatory mechanism is disrupted in plants lacking MCA channels **(Ma et al., 2025)**. Ethylene has also been implicated in responses to soil-compaction stress, where it mediates epidermal cell-wall thickening **(Zhang et al., 2025)**. Future experiments could therefore assess whether long-distance electrical communication activates ethylene signalling and leads to cell wall thickening in distal roots. Such a mechanism would be physiologically plausible: a root under attack could warn neighbouring roots to fortify their cell walls in preparation for potential biotic or abiotic challenges.

## Materials and methods

### Growth conditions and plant material

*Arabidopsis thaliana* seeds were sterilized with 70% ethanol for 5 minutes and 6% sodium hypochlorite (NaClO) + Tween 20 for 10 minutes. After six washing steps with H_2_O, seeds were sewn on Agar plates (1.5% Agar, 1.5% Sucrose, ½ MS, pH = 5.8), and stored at 4°C for at least two days. Aftrewards, plates were placed in a vertical position in a growth chamber (21 °C; 120 µmol photons.m^−2^.s^−1^) for 10-14 days. Plants expressing channelrhodopsins were grown under red light. The day before the experiment, seedlings were glued on Petri dish lids using medical adhesive (Medical Adhesive B Liquid, VM 355-1, Ulrich AG), and bathed in recording solution containing 1 mM CaCl_2_, 1 mM KCl, and 5mM of a MES/BTP pH buffer balanced at pH 6. On the next day, the plants were used for electrophysiology or imaging experiments.

The osca1 mutant seeds were kindly provided by Jian-Kang Zhu and Pengcheng Wang **(Lin et al., 2020)**, 35S-GCaMP3 seeds by Simon Gilroy **(Toyota et al., 2018)**, mca1 mutant seeds by Hubert Bauer **(Nakagawa et al., 2007)**, and the glr3.3/3.6 mutant seeds by Edward Farmer **(Mousavi et al., 2013)**. The retinal, ChR2 XXM2.0, and GtACR1 expressing plants were described in **(Song et al., 2025)**.

### Electrophysiology

Glued seedlings were placed on a microscope table (Akioskop 2FS, Zeiss, Germany), and illuminated with white light. To avoid photostimulation during optogenetic experiments, seedlings were illuminated with red light through a red-light glass filter with a cut-off wavelength of 635 nm. Seedling roots were observed through a water immersion objective (W Plan-Apochromat, 40x/1.0, Zeiss). A capillary filled with 300 mM KCl and sealed with 2% agarose in 300 mM KCl was placed in the bath solution, serving as a reference electrode. This reference electrode was connected to the ground via an Ag/AgCl half-cell.

Epidermal cells of the lateral root elongation zone were impaled with microelectrodes that were made from borosilicate glass capillaries (inner/outer diameter = 0.56/1.0 mm; Hilgenberg, Germany), which were pulled using a horizontal laser puller (P2000, Sutter Instruments, CA, USA). The electrodes were filled with 300 mM KCl, mounted on the holder of a piezo-driven micromanipulator (MM3A, Kleindiek, Reutlingen, Germany), and connected to a headstage via an Ag/AgCl half-cell. The headstage was linked to a custom-made microelectrode amplifier (Ulliclamp 1). Electrical signals were low-pass filtered at 0.5 kHz with a dual low-pass Bessel Filter (LPF 202A; Warner Instruments Corp., USA) and sampled at 1 kHz, while protocols were controlled by the WinWCP software **(Dempster, 1997)**. One or several pictures were taken following the recording in order to reconstruct the root network and calculate the distance between the recording electrode and the wounding capillary.

### Live fluorescence imaging and photostimulations

Arabidopsis seedlings expressing 35S-GCaMP3 were excited with an LED illumination system (pE-4000; CoolLED, Andover, UK) at 470 nm and filtered at 472±30 nm. The emission signals were passed through a dichroic mirror with a cut-off wavelength of 490 nm and a band filter at 520±35 nm. All filters were obtained from Semrock (BrightLine). Images were captured using a charge-multiplying charge-coupled device camera (QuantEM; Photometrics), and the whole setup was controlled by the Visiview software (Visitron). Recordings were performed at a 0.2 Hz sampling frequency. For optogenetic experiments, plants were stimulated with light pulses of 470nm during 1s every 30s with an intensity of 150 µmol.m^-2^.s^-1^. The retinal, ChR2 XXM2.0, and GtACR1 expressing plants were described in **(Song et al., 2025)**.

#### Wounding and local application

Glass capillaries were pulled using a horizontal puller (P2000, Sutter Instruments, CA, USA) and mounted on a micromanipulator. Pipettes with a tip size of ∼50 nm were used for single-cell wounding. For large wounds, pipette tips were broken to ∼10 µm, and capillaries were moved diagonally through at least 50% of the root cross-section. For local applications, pipette tips were broken to ∼10 µm, and a pressure of 2 bars was back-applied with the PDES 02DX (NPI Electronics) for 1s for sorbitol puffs, and 10s for other chemicals. A 10s puff corresponds to an ejection of 400 nL of intrapipette solution.

#### Analysis

All data were analysed with homemade Python scripts available in the following repository: https://github.com/AngelBaudon/Lateral-Root-Communication-2026.

Electrophysiology data were smoothed using a Gaussian filter (sigma = 10) and downsampled to 10Hz. The 10s before the stimulation were used as baseline and its mean were subtracted to the trace to calculate the depolarization amplitude. The rise and decay phases were fitted with a single exponential function and constants were measured as the time necessary to reach 63.2% or 36.8% of the depolarization, respectively. A single exponential decay was used in Figure 1 to correlate the depolarization amplitude with the distance between the wounding point and the recording site. This fitted model was then used to predict the depolarization in long-distance wounding experiments, and the measured depolarization was normalized by the predicted depolarization.

Imaging data were pre-processed using ImageJ to measure the background noise in a root-free area and the fluorescence variation at different hand-drawn ROI along the root. The background fluorescence values were subtracted to all other values. The 60s before the stimulation were used as baseline and its mean were subtracted to the trace and used to normalize this difference (ΔF/F_0_). Data were then smoothed using a Savitsky-Golay filter and the maximum Ca^2+^ signal was detected during the 60s following the stimulation.

Measurements were then exported to GraphPad Prism 8.0 for statistical analysis. Normality was tested before each test and a parametric test or its non-parametric equivalent was performed according to data distribution.

## Acknowledgments

This work was supported by the National Natural Science Foundation of China (W2531024) to RH, and by the Walter Benjamin program of the Deutsche Forschungsgemeinschaft (564693890) to AB.

## Author contributions

Project conception: AB, RH; Methodology: AB, SH, RR; Experimentation: AB; Writing, AB, RH, RR, DG, DB, SH; Project administration and supervision: AB, RH, DG.

## Supplementary information

**Supplementary Figure 1.**
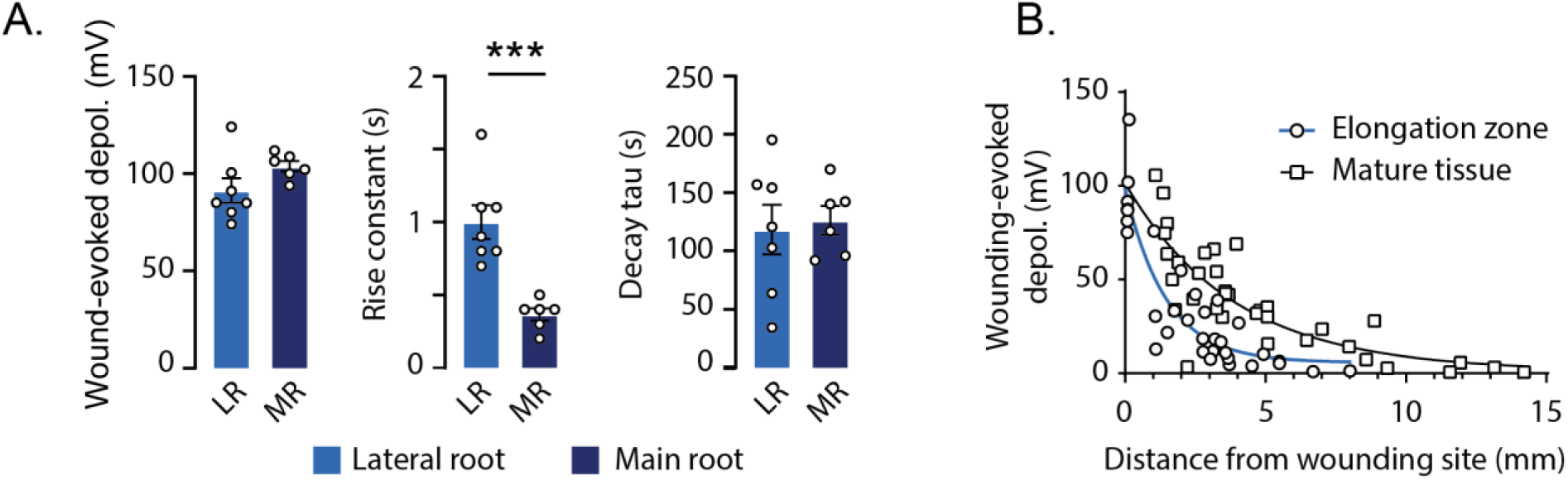
**A.** Amplitude, rise, and decay constants of the depolarization triggered by a large wounding of a lateral root (LR), or of the main root (MR). **B.** Amplitude of the wounding-evoked depolarization plotted as a function of the distance between the wounded root and the recording point located in mature root tissues (squares), or in the elongation zone of another lateral root (circles). Each point is from an individual wounding experiments. Data are expressed as mean across plants ± SEM. Detailed statistics can be found in the *Statistics Table*. ***p<0.001

**Supplementary Figure 2.**
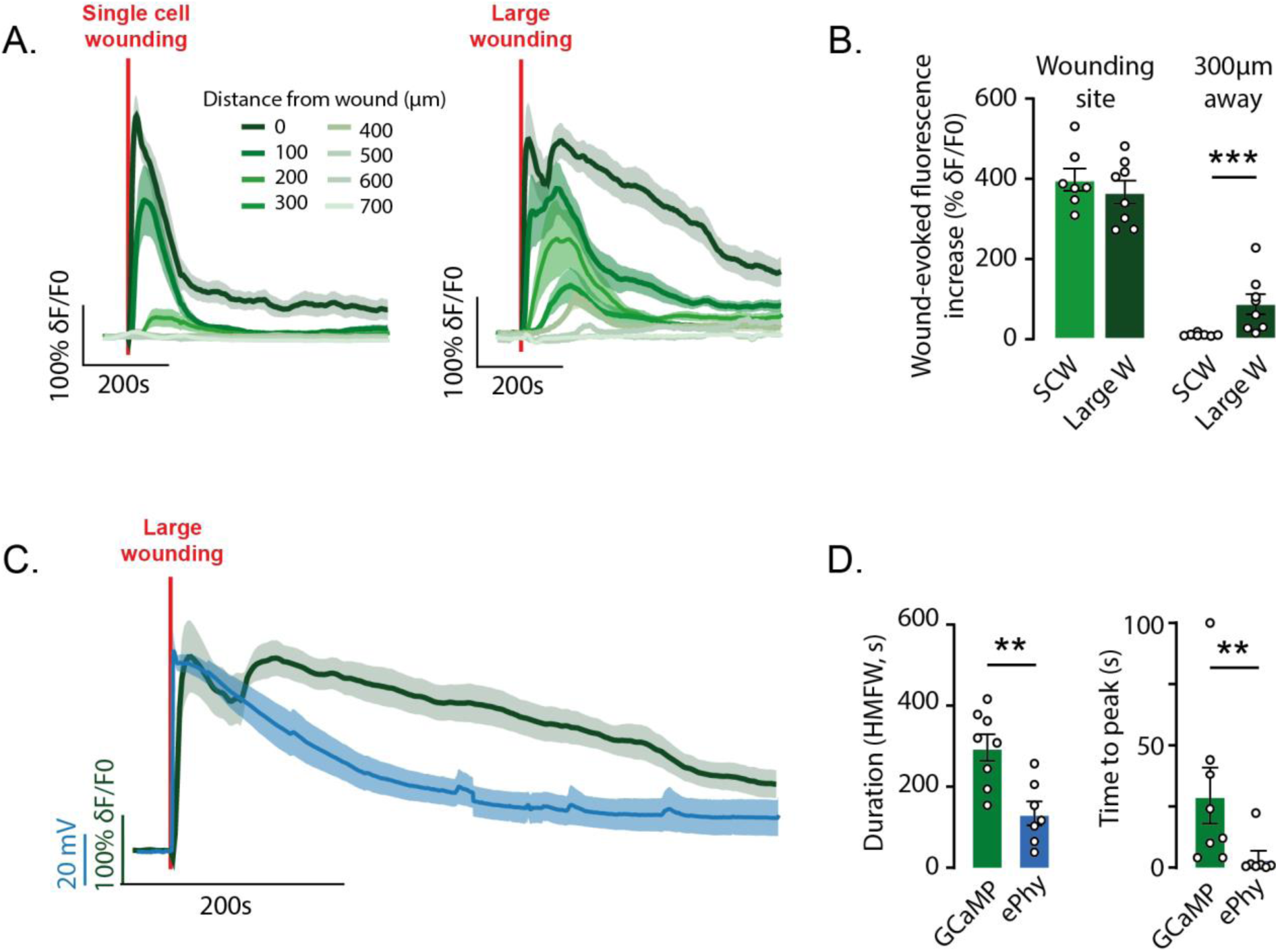
**A.** Average fluorescence levels of GCaMP3 measured at several distances from the site of a single cell wounding (left, n=7), or a large wounding (right, n=8). Each trace is from an individual wounding experiments. The red line indicates the time point of wounding. **B.** Quantification of GCaMP3 fluorescence at the wound site and 300µm away following single-cell (light green) or large wounding (dark green). **C.** Overlay of epidermal cell membrane potential (blue), and cytosolic Ca^2+^ level (green) levels following wounding. Each trace is from an individual wounding experiments. The red line indicates the time point of wounding. **D.** Duration measured as the half-maximum full width (HMFW, in seconds), and time to peak of the data shown in C. Data are expressed as mean across plants ± SEM. Detailed statistics can be found in the *Statistics Table*. **p<0.01, ***p<0.001.

**Supplementary Figure 3.**
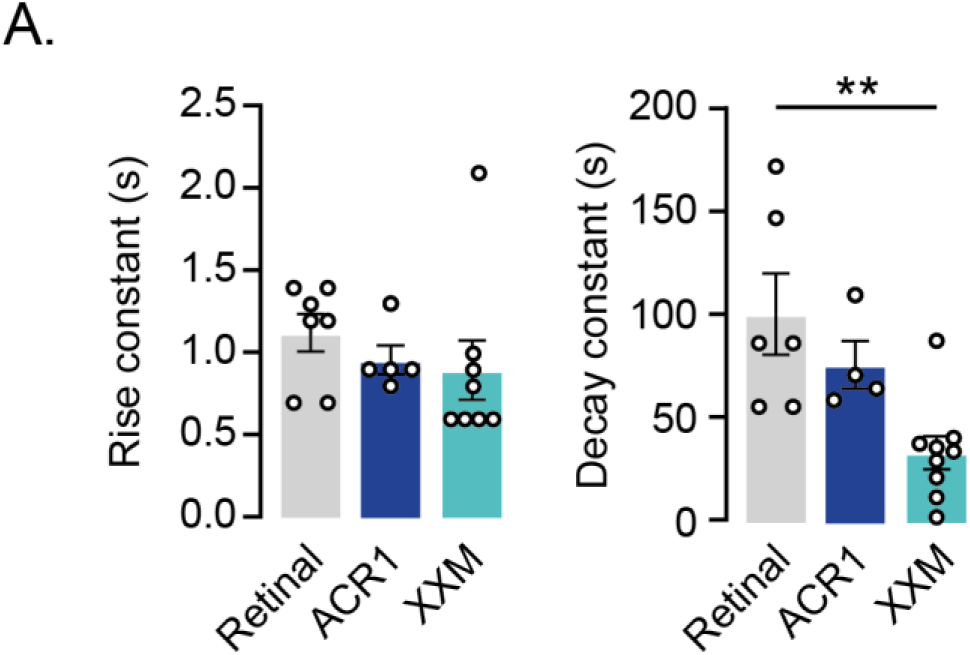
**A.** Rise and decay constant of the wound-evoked depolarization following repetitive optogenetic stimulation. n_Retinal_=5, n_GtACR1_=9, n_XXM_=9. Each point is from an individual wounding experiments. Data are expressed as mean across plants ± SEM. Detailed statistics can be found in the *Statistics Table*. **p<0.01.

**Supplementary Figure 4.**
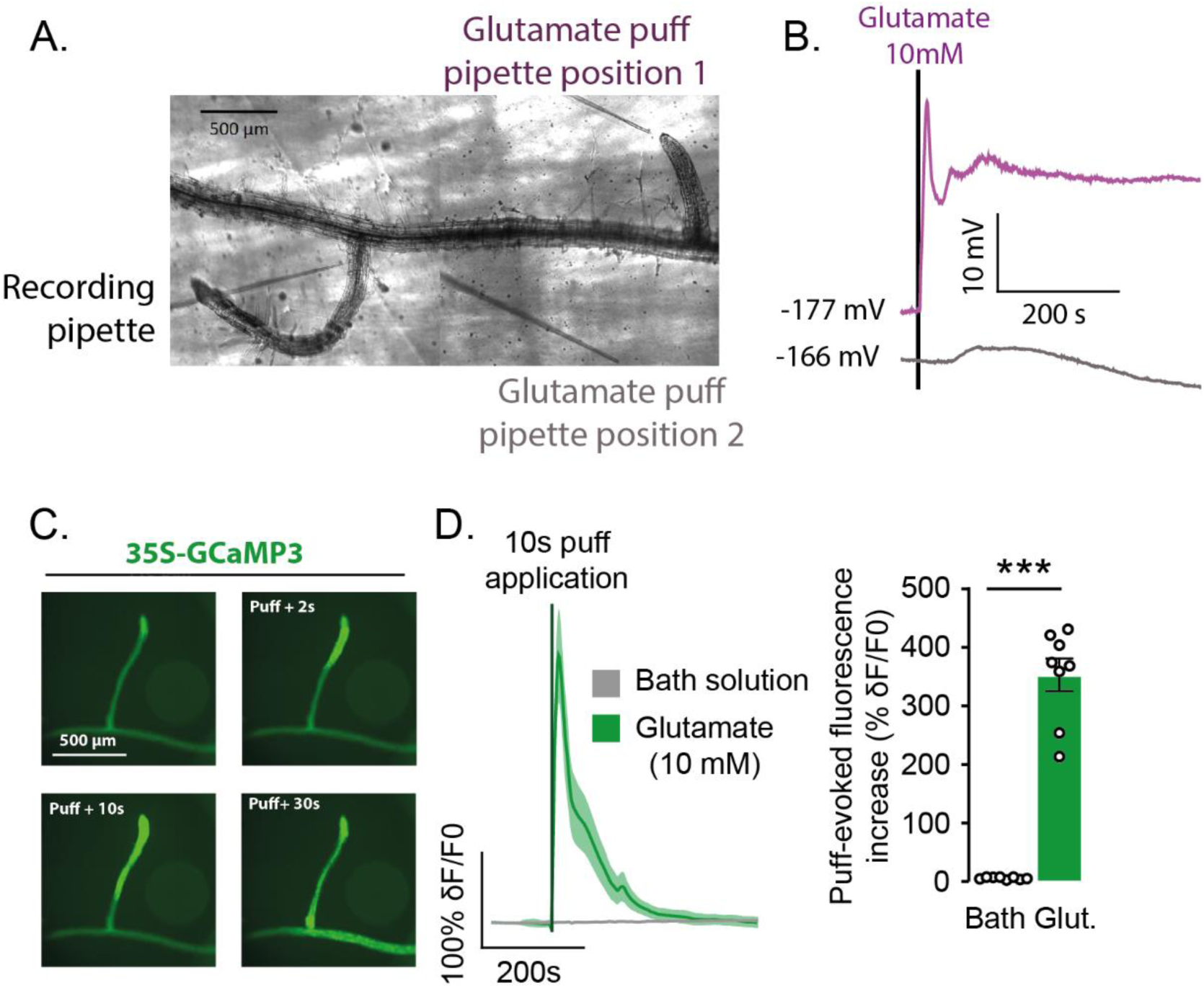
**A-B.** Control of local glutamate diffusion in the bath. **A.** Micrograph showing the position of the recording pipette, the puff application pipette on a distant root (position 1), and on a place without any root (position 2). **B.** Membrane potential recorded during local glutamate application on position 1 or in position 2 shown in panel A. The black line indicate the glutamate application time point. **C.** Fluorescence imaging of a lateral root expressing GCaMP3 at different time points before, during, and after a puff application of glutamate (10mM). Scale = 500µm. **D.** Mean GCaMP3 fluorescence variation (left), and corresponding quantification (right).The black line indicate the glutamate application time point. Each trace is from an individual wounding experiments. Data are expressed as mean across plants ± SEM. Detailed statistics can be found in the *Statistics Table*. ***p<0.001.

**Supplementary Figure 5.**
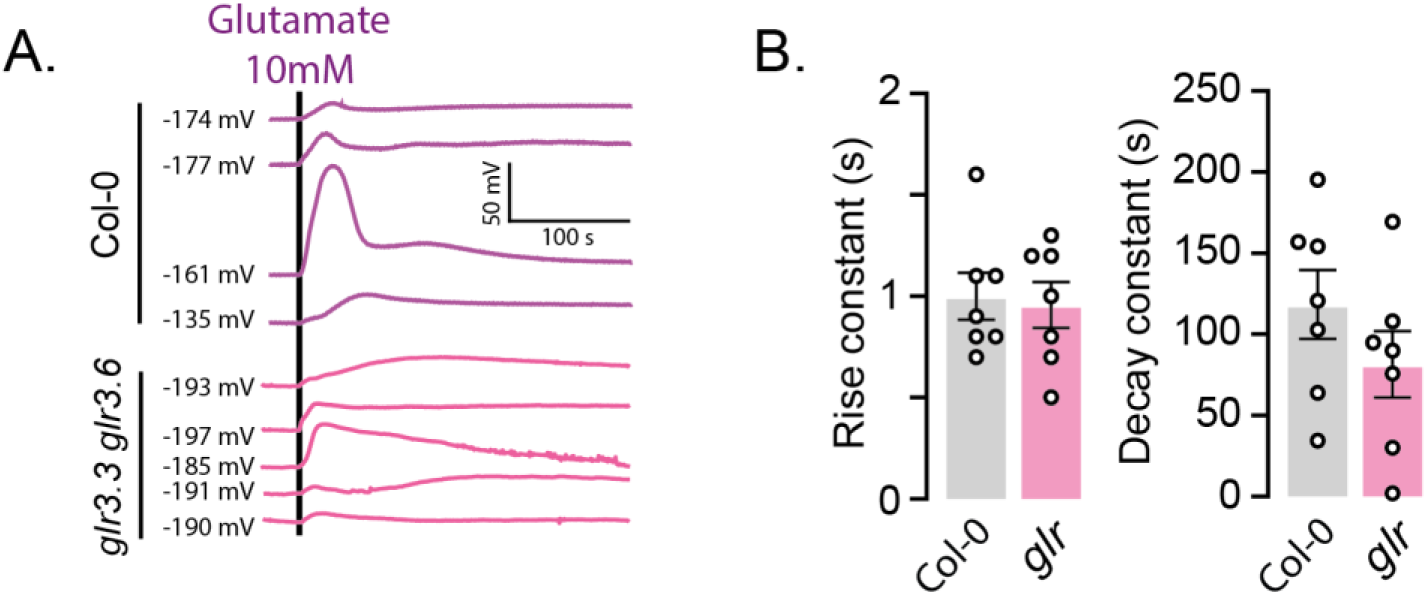
**A.** Membrane potential depolarization triggered by distant glutamate application in wild type and *glr3.3/3.6* mutant plants. Each trace is from an individual wounding experiments. The black line indicate the glutamate application time point. **B.** Rise and decay constant of wound-evoked depolarization recorded in *glr3.3/3.6* mutant and wild type plants. Data are expressed as mean across plants ± SEM. Detailed statistics can be found in the *Statistics Table*.

**Supplementary Figure 6.**
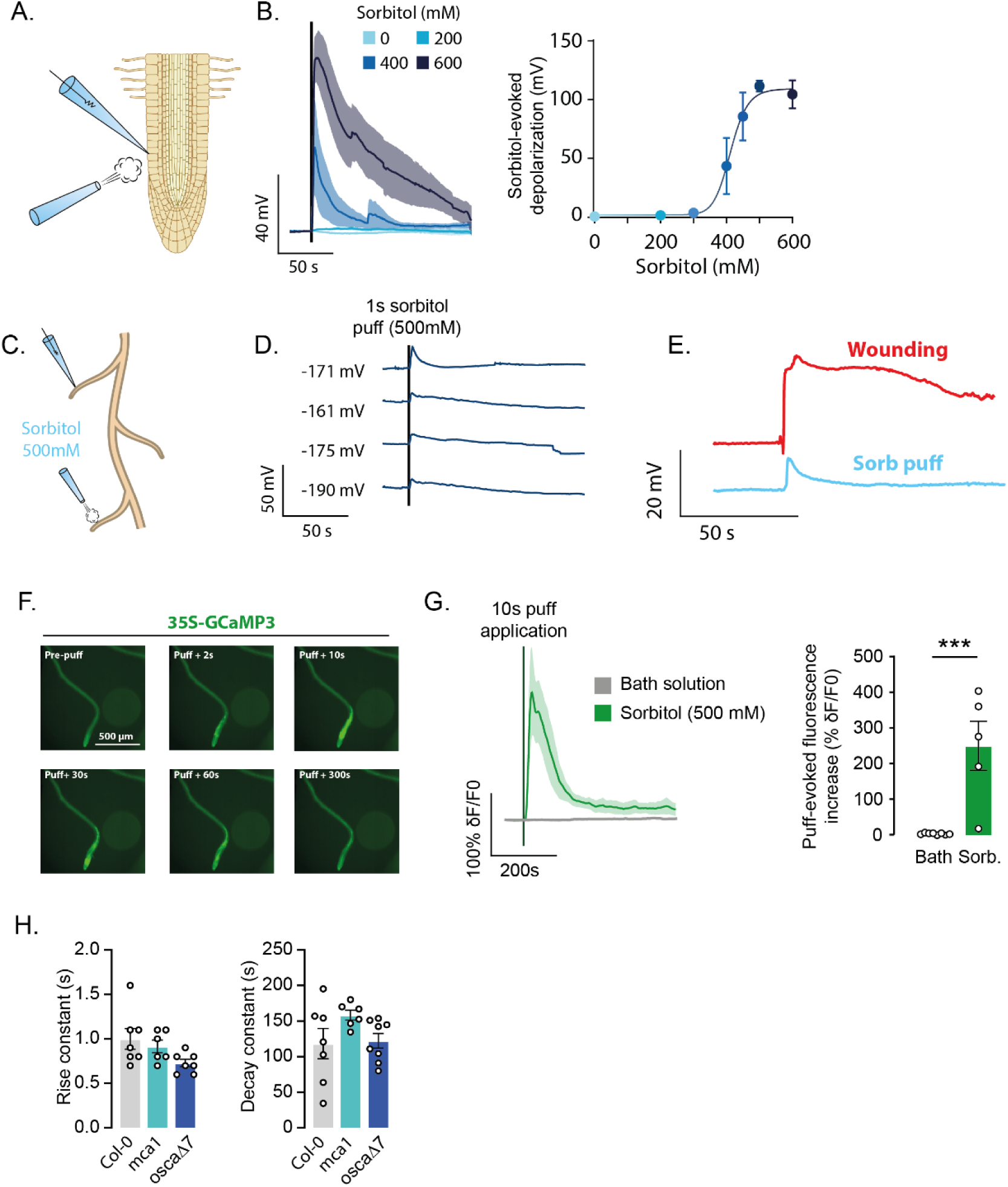
**A.** Epidermal root cells were impaled, and a solution containing different concentrations of sorbitol was backpressure-applied locally with a second glass pipette for 1s. **B.** Mean traces of sorbitol-evoked depolarization (left), and corresponding quantification (right). Sorbitol application time point is indicated with the black line. Each trace come from independent experiments. **C.** The epidermal cell of a lateral root was impaled using a sharp electrode, and sorbitol was locally applied on another, distant lateral root. **D.** Example traces of membrane potential depolarization evoked by sorbitol application on another distant lateral root. Each trace comes from a different plant; the number before the trace indicates the membran2e potential at the beginning of the experiment, and the red line indicates the wounding time point. **E.** Comparison of the depolarizations evoked by applying sorbitol or wounding a distant root. The two recordings were performed in the same pair of root. **F.** Fluorescence imaging of a lateral root expressing GCaMP3 at different time points before, during, and after a puff application of sorbitol (500mM). **G.** Mean GCaMP3 fluorescence variation (left), and corresponding quantification (right). Each data point comes from a different plants. The black line indicate the sorbitol application time point. **H.** Rise and decay constant of wound-evoked depolarization recorded in plants lacking functional mechanosensitive channels mca1, osca1, and in their controls (Col-0). Each data point comes from a different plants. Data are expressed as mean across plants ± SEM. Detailed statistics can be found in the *Statistics Table*. ***p<0.001.

